# Lytic bacteriophages against *Salmonella* Typhi as a potential alternative to antibiotics

**DOI:** 10.64898/2026.07.11.737897

**Authors:** Sujit Tandukar, Padma Shrestha, Manisha Shrestha, Basudha Shrestha, Anjana Singh, Reshma Tuladhar, Jivan Shakya

## Abstract

**Introduction:** Enteric fever, being endemic with seasonal peaks in low- and middle-income countries, is a major health concern. Moreover, the rise in antibiotic resistance has exacerbated the situation. This study was undertaken to investigate the lytic bacteriophages against *Salmonella* Typhi with a potential for phage therapy.

**Materials and Methods:** A hospital-based cross-sectional study was conducted from October 2023 to March 2024. Blood cultures were processed by the BACTEC automated culture system following standard microbiological techniques to isolate typhoidal *Salmonella*. Antibiotic susceptibility was tested by the modified Kirby-Bauer disc diffusion method. Lytic bacteriophages isolated by the double-layer agar method were assessed for their host range and lytic ability with spot and turbidimetric assays.

**Results:** Of the total 1054 blood specimens, 35 (3.2%) were positive for *S. Typhi*. All the isolates were susceptible to first-line antibiotics—ampicillin, chloramphenicol, and cotrimoxazole. The isolates were also sensitive to nalidixic acid (80%) as well as fluoroquinolones; ciprofloxacin (62.86%), levofloxacin (77.14%), and ofloxacin (80%). Fifteen lytic phages were isolated against *S. Typhi* Ty2 and CT18 strains. Four phages—vB_SaTy_ST2, vB_SaTy_ST7, vB_SaTy_ST17, and vB_SaTy_ST18—lysed all 35 clinical *S.* Typhi isolates. While vB_SaTy_ST17 and vB_SaTy_ST18 also lysed 7 out of 20 *S.* Paratyphi A isolates. Three phages (vB_SaTy_ST2, vB_SaTy_ST7, vB_SaTy_ST17) were tested against *S.* Typhi isolate S30. Individually, vB_SaTy_ST17 suppressed the growth for 13 hours, vB_SaTy_ST2 and vB_SaTy_ST7 for 10 hours. The phage cocktail vB_SaTy_ST2 + vB_SaTy_ST17 was the most effective, which extended the inhibition time to 15 hours.

**Conclusion:** This study highlights the ongoing burden of enteric fever in Nepal and the increase in susceptibility of *S*. Typhi to nalidixic acid and fluoroquinolones. It also demonstrates the promising lytic potential of bacteriophages, particularly vB_SaTy_ST17 and the phage cocktail vB_SaTy_ST2 + vB_SaTy_ST17, against clinical *S.* Typhi, highlighting their potential as alternatives to antibiotics.

## Introduction

Enteric fever is an acute and often life-threatening febrile illness caused by the Gram-negative bacterium *Salmonella enterica* subspecies *enterica* serovar Typhi (*S.* Typhi) and *Salmonella enterica* serovar Paratyphi (*S.* Paratyphi). The predominant manifestation of typhoid fever is sustained high fever with other nonspecific symptoms like chills, nausea, and abdominal pain [1,2]. Globally, 14.3 million cases of typhoid and paratyphoid fevers occurred in 2017, with the majority (76·3%) of infections by *S.* Typhi. The case fatality is high in the children, aging population, and those living in lower-income countries [3]. Case fatality rate of typhoid fever significantly reduced to 1-4% from 10-30%, with implementation of proper antibiotic therapy [1].

Resistance to first-line drugs for enteric fever such as, chloramphenicol, ampicillin, and cotrimoxazole is considered multidrug resistance (MDR), thus fluoroquinolones, third-generation cephalosporins, and azithromycin are the alternatives [4,5]. The first extensively drug-resistant (XDR) *S*. Typhi reported from Pakistan in 2016 [6], which spread to the Middle East and further to North America, Europe, and Australia, was resistant to fluoroquinolone and third-generation cephalosporin, besides the first-line drugs [6,7]. The only effective antimicrobials left for XDR *S.* Typhi are azithromycin (oral antibiotic) and carbapenems (intravenous antibiotic) [7]. However, sporadic cases of azithromycin-resistant *S*. Typhi recently reported from Bangladesh, Pakistan, Nepal, and India have challenged the treatment of enteric [8–10] necessitating an effective alternative.

Revival of the phage therapy, a century-old method dating back to 1919 when Félix d’Hérelle used phages to treat children with bacterial dysentery [11] is one of the promising approach to address the growing antibiotic resistance [12]. Bacteriophages (or simply phage), the viruses that infect bacteria abundantly found in the environment with 10-fold greater than bacteria [13,14]. Bacteriophages recognize their host via highly specific receptors, ensuring they are precise for the target pathogen and have minimal or no effect on the normal flora, making them an ideal therapeutic agent [15].

Thus, this study aims to investigate the bacteriophages isolated from wastewater samples that exhibit lytic activity against *S.* Typhi.

## Materials and Methods

### Study site, design, and period

A cross-sectional study was conducted at Kathmandu Model Hospital, Kathmandu, Nepal, and the Institute for Research in Science and Technology (IRSTech), Lalitpur, during the period of six months from October 2023 to March 2024.

### Study population and sampling technique

The blood samples of the patients requested for blood culture by the clinician visiting Kathmandu Model Hospital during the study period were included in the study through a convenience sampling technique.

### Blood collection and processing

Blood specimens were aseptically collected by experienced laboratory staff using vein puncture. For adults, 10 mL of blood was collected, while 2 to 3 mL for paediatric patients, and immediately transferred to BACTEC culture bottles, then incubated at 37°C for 5 days using the BACTEC automated culture system (Becton Dickinson, Franklin Lakes, NJ, USA). The positive samples were sub-cultured in MacConkey and Blood agar and incubated at 37⁰C for 18 hours. The bacterial species were identified by standard microbiological procedures following colony morphology, Gram staining, and biochemical tests [16].

### Antibiotic susceptibility testing

Antimicrobial susceptibility testing (AST) was performed on Muller-Hinton Agar using the Kirby-Bauer Disc Diffusion technique according to the Clinical and Laboratory Standards Institute (CLSI) guidelines [17]. The antibiotic discs used were amoxicillin (10 μg), chloramphenicol (30 μg), cotrimoxazole (25 μg), nalidixic acid (30 μg), ciprofloxacin (5 μg), ofloxacin (5 μg), levofloxacin (5 μg), cefixime (5 μg), cefotaxime (30 μg), ceftriaxone (30 μg), gentamycin (10 μg), and azithromycin (15 μg). After incubation, the diameters of zones of inhibition around the discs were measured, and the results were interpreted as sensitive, intermediate, or resistant, using an interpretative criterion of the CLSI.

### Bacteriophage isolation and purification

Bacteriophages were isolated from the sewage and river water by the double-layer agar (DLA) method [18] with some modifications using *S.* Typhi Ty2 and *S.* Typhi CT18 as hosts. Briefly, the water sample was centrifuged at 4500 rpm for 10 minutes to remove debris and filtered through a 0.22 μm membrane filter. One millilitre of filtrate was mixed with 0.5 ml of the exponential phase of the host strain and incubated at room temperature for 10 minutes, to which 3 ml of molten soft agar (0.75%) was added. The content was then overlaid over a solid bottom nutrient agar (NA) plate and incubated for 24 h at 37°C. The plaques were selected based on morphology, and diameter. For purification, a typical plaque was picked from the top layer using a sterile pipette tip, suspended in 1 ml SM buffer (50 mM Tris–HCl, 10 mM NaCl, 8 mM MgSO_4_, pH 7.4), and vortexed. Phage was purified by the DLA method as described previously. The purification process was performed repeatedly three times. Phages were suspended in SM buffer and stored at 4°C until further use.

### Determination of host range

The host range of isolated phages was determined by spot test with some modifications [19]. Briefly, a 5 μl phage stock solution was spotted on the lawn of different bacterial strains and incubated at 37°C overnight. The plates were observed for the presence of clear lytic zones. The bacterial strains tested were *E. coli* ATCC 25922, *S. aureus* ATCC 29213, *S. aureus* ATCC 43300, *E. faecalis* ATCC 51299, *E. faecalis* ATCC 29212, *Klebsiella quasipneumoniae* ATCC 700603, *Klebsiella pneumoniae* BAA-1705, *Pseudomonas aeruginosa* ATCC 27853, as well as four *S.* Typhi strains- *S.* Typhi Ty2, *S*. Typhi CT18, *S.* Typhimurium LT12, and *S*. Typhi ΔVi (Vi capsule knock-out mutant).

### Lytic ability against clinical isolates

The lytic ability of isolated phages against clinical *Salmonella* isolates was determined by the spot test described previously [19]. Briefly, a 5 μl phage stock solution was spotted on the lawn of the clinical isolate and incubated at 37°C overnight. The phage exhibiting clear zones on the spotted region was considered to have lytic activity, while those that did not produce clear zones or produced turbid zones were considered to have no lytic activity against the tested organism. The clinical *S*. Typhi isolates for the lytic assay were obtained during this study period, while the clinical *S*. Paratyphi isolates were obtained from the microbiology laboratory of Kathmandu Model Hospital.

### In-vitro efficacy of phage and phage cocktail against clinical *S.* Typhi isolate

The lytic ability of phages and their cocktail was evaluated against fluoroquinolone-resistant *Salmonella* Typhi isolate S30 *in vitro* by a turbidimetric growth kinetic assay described elsewhere, with some modifications [20]. Briefly, the overnight culture of the bacterium was diluted in fresh tryptone soya broth (TSB) medium to reach the final OD_600_ of ∼ 0.2. Three phages, vB_SaTy_ST2, vB_SaTy_ST7, and vB_SaTy_ST17, were selected based on their broad lytic ability. The bacterium (10^8^ CFU/mL) was infected with respective phages (10^7^ PFU/mL) to the final MOI of 0.1 and incubated at 37°C for 24 hours. The bacterial growth was determined by measuring the OD_600_ at intervals of one hour using a spectrophotometer. Sterile TSB medium was used as a blank for the measurement, while S30 without phage infection was used as the positive control. A standard curve of time versus OD_600_ readings was plotted, with the results presented as the average of triplicate. The OD_600_ reading of 0.1 was considered an indicator of early log phage.

For evaluating the efficacy of the phage cocktail, respective phages were mixed in equal proportion to the final concentration of 10^7^ PFU/mL and mixed with S30 at a concentration of 10^8^ CFU/mL to achieve an MOI of 0.1, followed by incubation at 37°C for 24 hours. The optical density readings were measured at the intervals of one hour, and the standard curve was plotted.

### Quality control

Culture media plate from each batch was incubated at 37°C for 24 hours to check for sterility. The culture media were visually inspected for microbial growth or deterioration before sample inoculation. *Escherichia coli* ATCC 25955 and *Staphylococcus aureus* ATCC 25923 standard strains were used as quality controls for the standardization of AST.

### Ethical approval

The study was approved by the Institutional Review Committee (IRC) of the Public Health Concern Trust, Nepal (phect- NEPAL), Kathmandu Model Hospital, Kathmandu (Regd. No: 124-2023). The formal informed consent was waived by the IRC of phect- Nepal.

### Data analysis

Data were recorded in Microsoft Excel and analysed using RStudio 2024.04.1+748 (R version 4.4.0) for Windows. Descriptive statistics were used to summarize the demographic characteristics of study participants, bacterial isolates, AST patterns, and bacteriophage lytic ability. Fischer’s exact test was used to assess the statistical significance of phage lytic ability. p < 0.05 was considered statistically significant.

## Results

### Prevalence of *S*. Typhi

During the study period, a total of 1054 blood specimens were processed for bacteriological culture. Among these specimens, 5.2% (55/1054) were positive for bacterial growth, out of which 3.32% (35/1054) were identified as *S.* Typhi (Table 1).

**Table 1.**
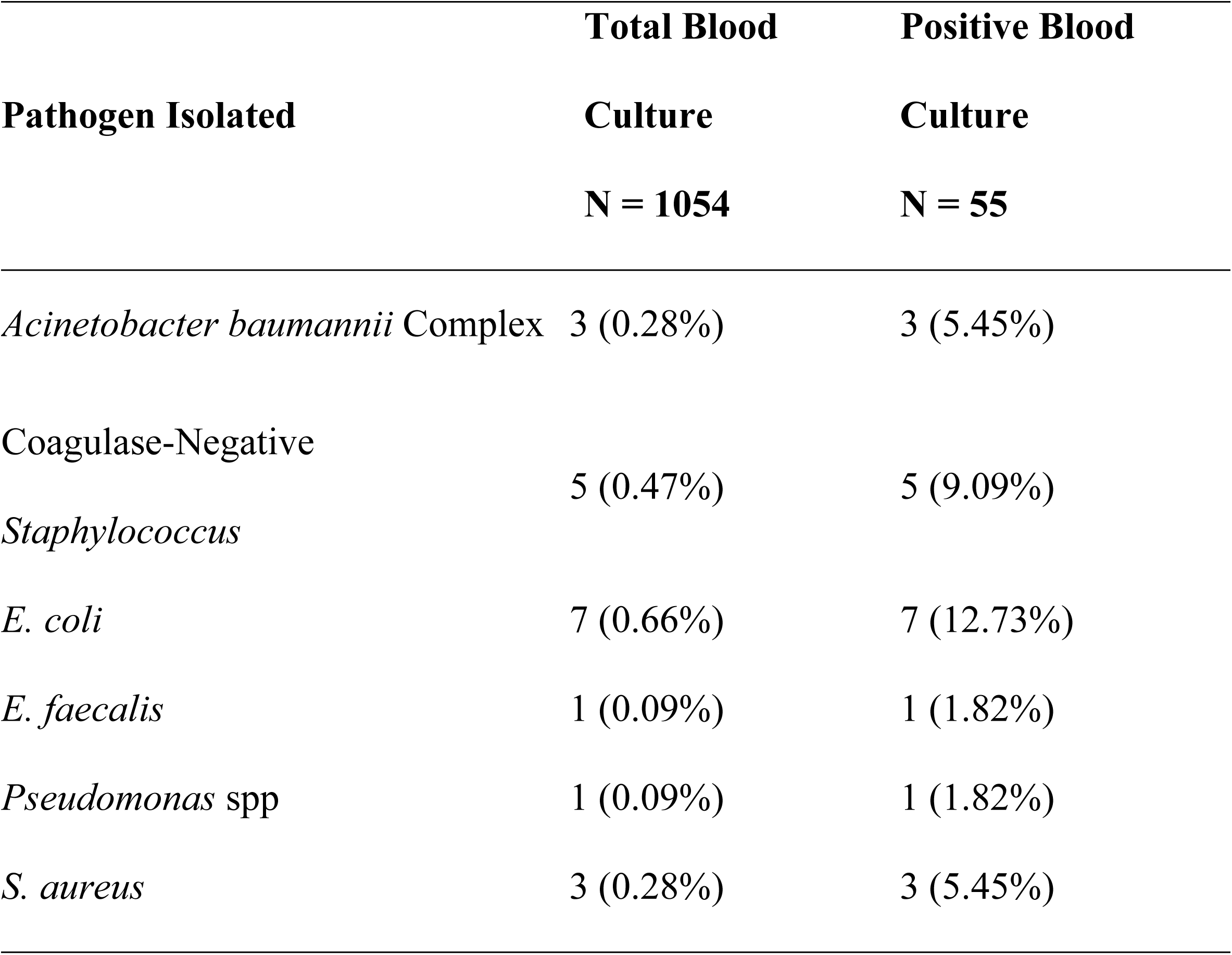
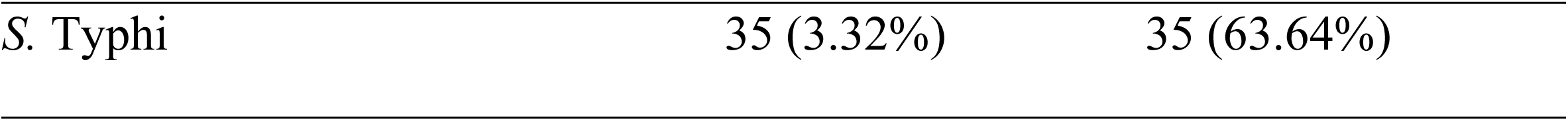
Distribution of bacterial isolates in the blood specimen.

### Distribution of *S*. Typhi by sociodemographic characteristics

Among thirty-five *S*. Typhi, 21 (60%) were isolated from male and 14 (40%) from female. Typhoid fever was more prevalent in the age group 16-25 years (48.57%) compared to other age groups, with the mean and median age of 24.4 years and 22 years, respectively. However, typhoid fever was not observed in the age groups <5 years and >56 years (Table 2).

**Table 2.**
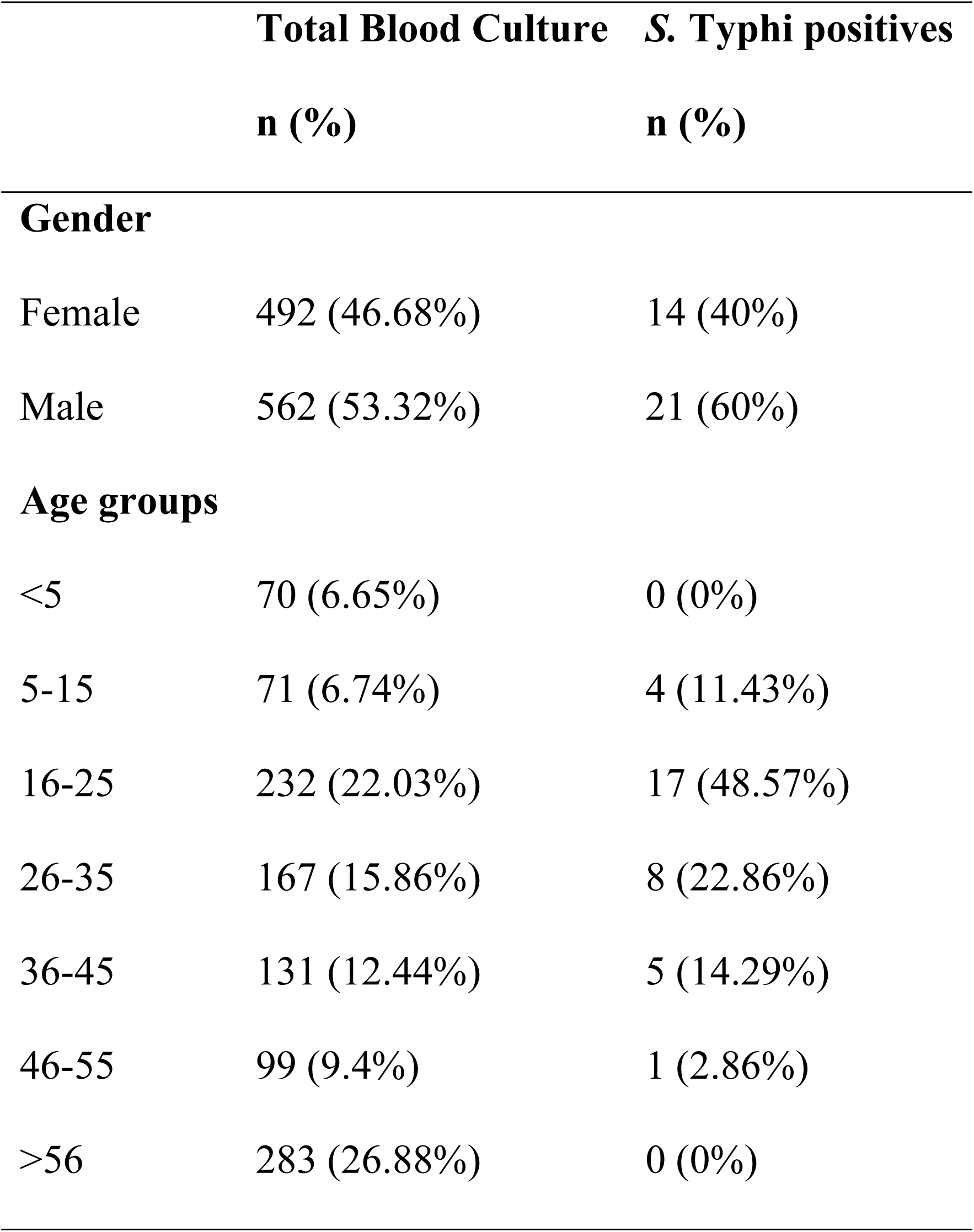
Socio-demographic distribution of study participants with *S.* Typhi infection.

|  | Total Blood Culture | *S. Typhi* positives |
| --- | --- | --- |
|  | n (%) | n (%) |
| **Gender** |  |  |
| Female | 492 (46.68%) | 14 (40%) |
| Male | 562 (53.32%) | 21 (60%) |
| **Age groups** |  |  |
| <5 | 70 (6.65%) | 0 (0%) |
| 5-15 | 71 (6.74%) | 4 (11.43%) |
| 16-25 | 232 (22.03%) | 17 (48.57%) |
| 26-35 | 167 (15.86%) | 8 (22.86%) |
| 36-45 | 131 (12.44%) | 5 (14.29%) |
| 46-55 | 99 (9.4%) | 1 (2.86%) |
| >56 | 283 (26.88%) | 0 (0%) |

### Distribution of *S*. Typhi in different months of the year

Most of the *S*. Typhi were isolated in October (68.57%), followed by November (14.29%). However, *S.* Typhi isolated from the blood samples every month from December to March was in a low proportion (Fig 1).

**Fig 1.**
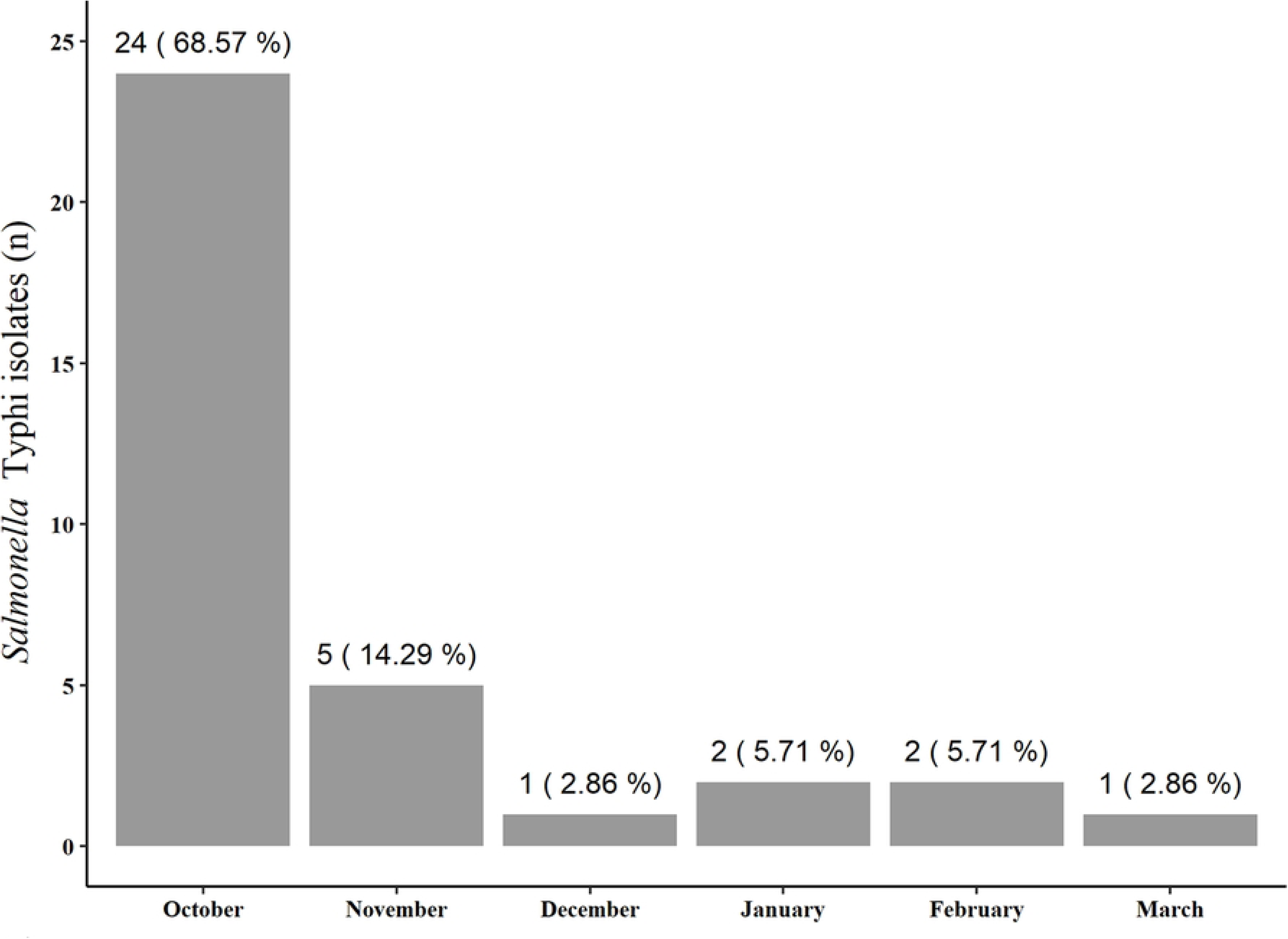
Month-wise distribution of *S*. Typhi isolates

### Antibiotic susceptibility pattern of isolates

All the *S*. Typhi isolates were susceptible to amoxicillin, azithromycin, chloramphenicol, cefixime, ceftriaxone, cefotaxime, gentamicin, and cotrimoxazole. The isolates were also sensitive to nalidixic acid (80%) as well as fluoroquinolones; ciprofloxacin (62.86%), levofloxacin (77.14%), and ofloxacin (80%) (Fig 2).

**Fig 2.**
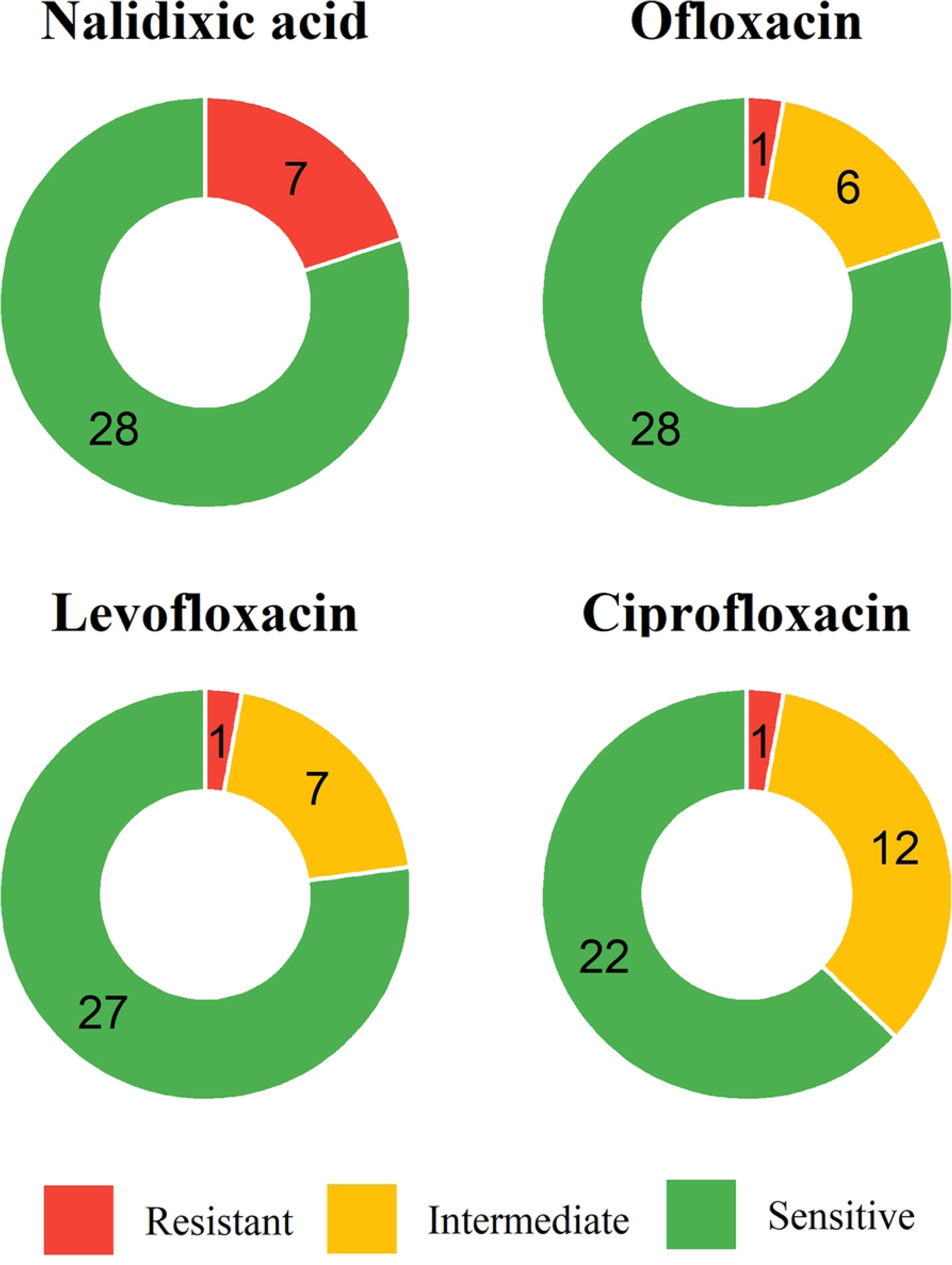
Antibiotic susceptibility pattern of *S.* Typhi against fluoroquinolones. AST was performed using the Kirby-Bauer Disc Diffusion technique, and the isolates were classified as sensitive, intermediate, or resistant following interpretation criteria from CLSI. The numbers within the chart indicate the actual isolate counts for each category.

### Bacteriophage isolation

A total of fifteen bacteriophages were isolated from 4 different water samples, which included river water (n = 2) and sewage water (n = 2). Among them, eleven (73.33%) phages were isolated from sewage and 4 (26.67%) from river water. (Figs 3 and 4; Table 3). Eleven out of the fifteen bacteriophages were isolated against the *S.* Typhi Ty2 strain, while four were isolated against the MDR *S.* Typhi CT18 strain. Bacteriophages infecting the Ty2 strain formed large-sized plaques (1 – 4 mm) while those infecting CT18 strains formed small plaques (pinpoint or pinhead) (S1 Table).

**Fig 3.**
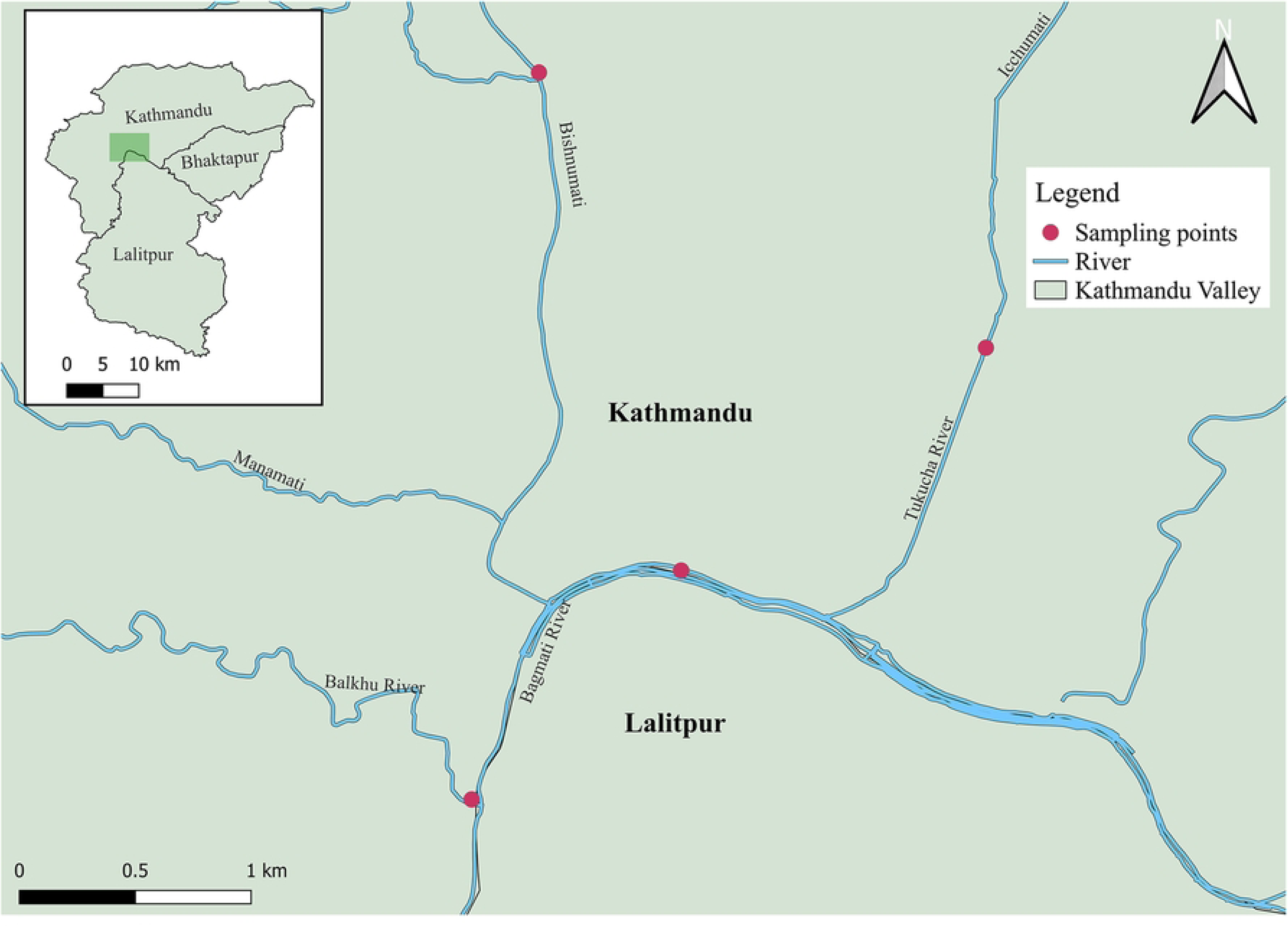
Sampling locations for isolation of bacteriophage in Kathmandu, Nepal. The base map is from OpenStreetMap, and the rendering was done using QGIS (version 3.40).

**Fig 4.**
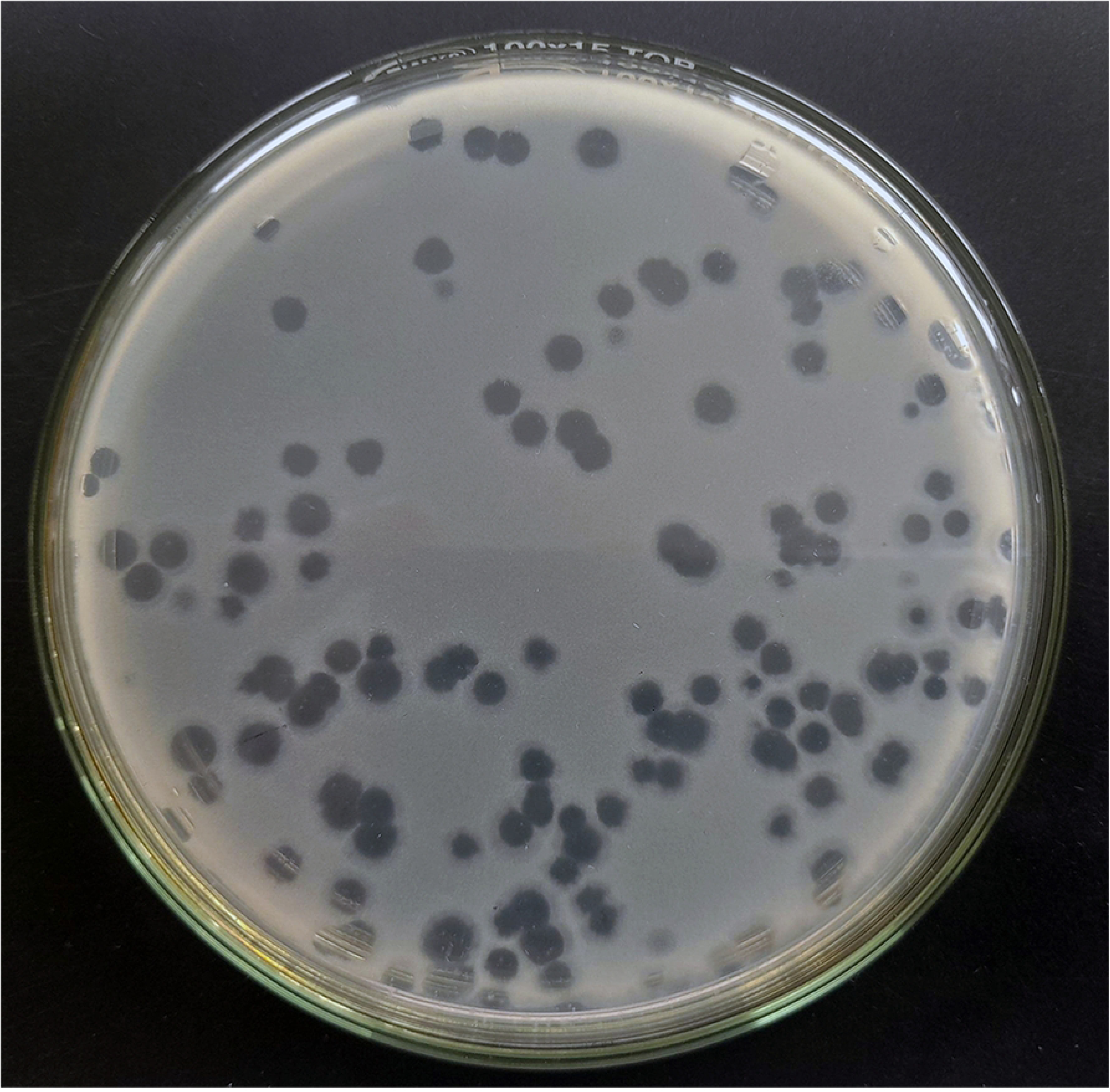
Isolation of bacteriophages against *S.* Typhi using the DLA method. One mililiter of sewage contaminated river water was mixed with 0.5ml of S. Typhi (OD600 = 0.2) and mixed with molten TSA (0.75%) and overlaid onto nutrient agar plate and incubated at 37°C for 24hrs. The clear spots indicate presence of lytic bacteriophages against S. Typhi.

**Table 3.**
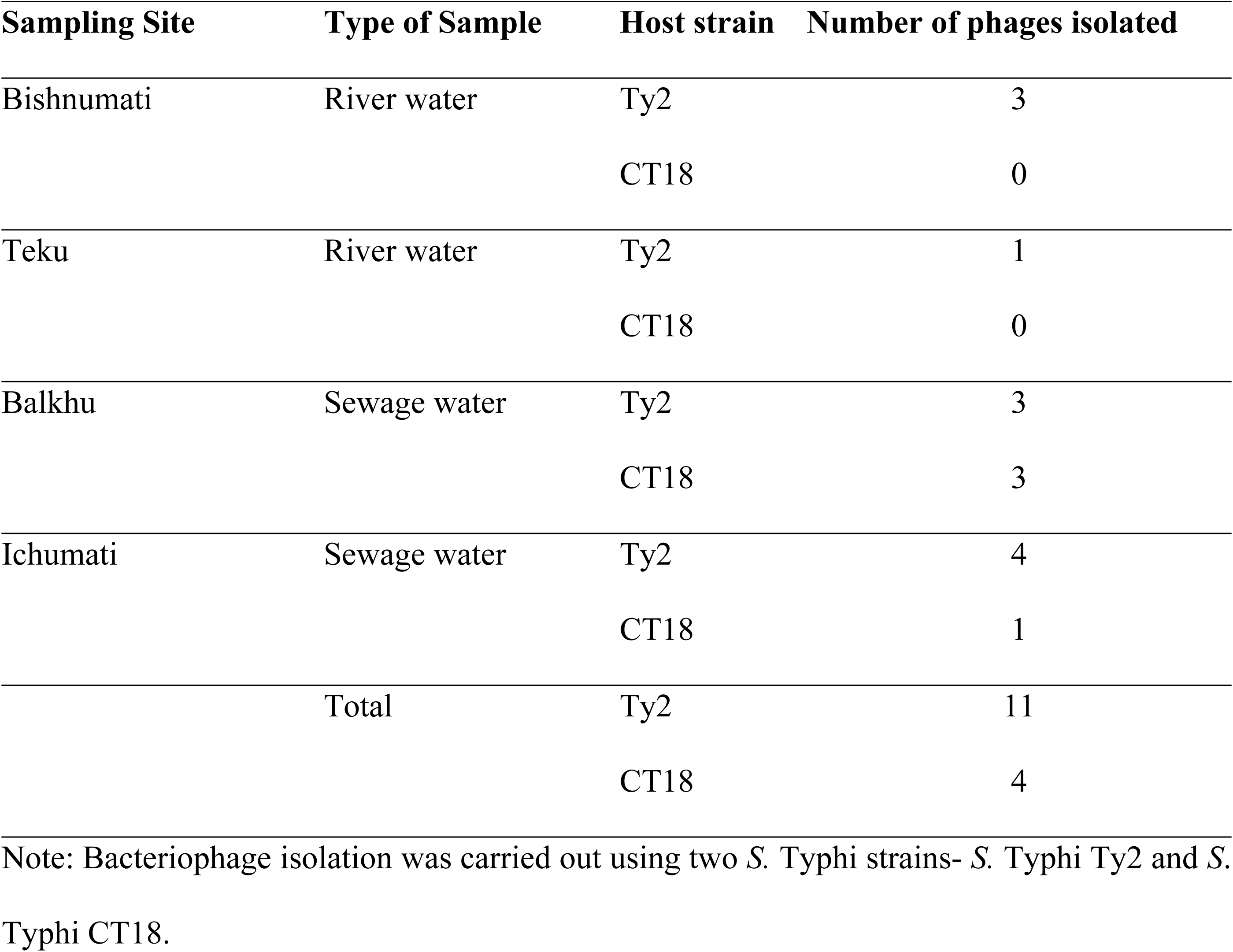
***S*. Typhi-specific phage isolation from environmental samples collected from**

| Sampling Site | Type of Sample | Host strain | Number of phages isolated |
| --- | --- | --- | --- |
| Bishnumati | River water | Ty2 | 3 |
|  |  | CT18 | 0 |
| Teku | River water | Ty2 | 1 |
|  |  | CT18 | 0 |
| Balkhu | Sewage water | Ty2 | 3 |
|  |  | CT18 | 3 |
| Ichumati | Sewage water | Ty2 | 4 |
|  |  | CT18 | 1 |
|  | Total | Ty2 | 11 |
|  |  | CT18 | 4 |

Note: Bacteriophage isolation was carried out using two *S. Typhi* strains- *S. Typhi* Ty2 and *S.* *Typhi* CT18.

### Host range

All isolated phages exhibited high specificity and could not infect bacteria from other genera, namely *E. coli* ATCC 25922, *S. aureus* ATCC 29213, *S. aureus* ATCC 43300, *E. faecalis* ATCC 51299, *E. faecalis* ATCC 29212, *Klebsiella quasipneumoniae* ATCC 700603, *Klebsiella pneumoniae* BAA-1705, and *Pseudomonas aeruginosa* ATCC 27853. Furthermore, the phages isolated against the *S*. Typhi Ty2 strain could not infect different strains within the same species- *S*. Typhi CT18 and *S.* Typhimurium LT12. A similar observation was seen in phages isolated against *S*. Typhi CT18. None of the pages were able to infect the Vi polysaccharide knock-out *S*. Typhi ΔVi strain (Table 4).

**Table 4.**
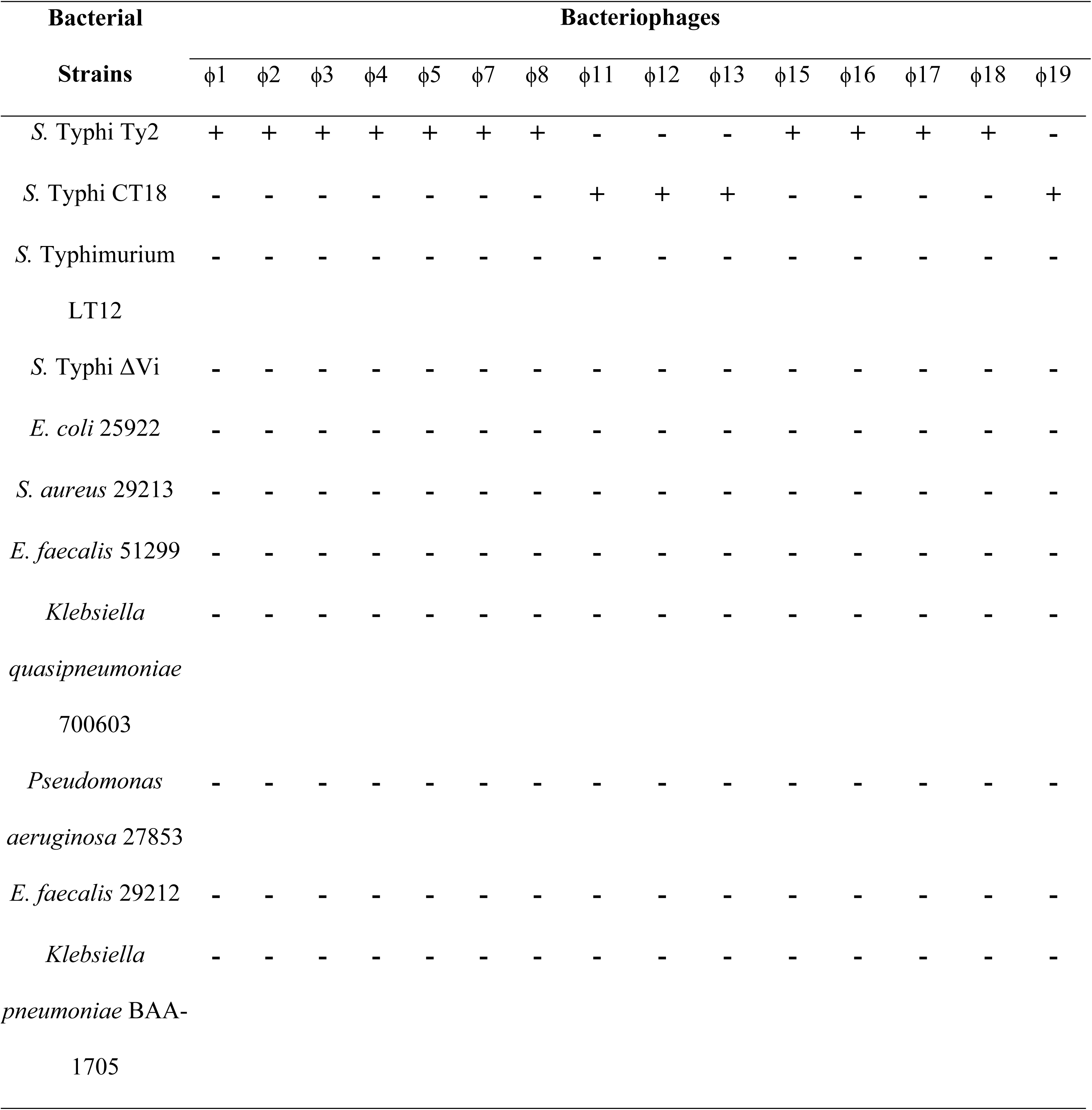
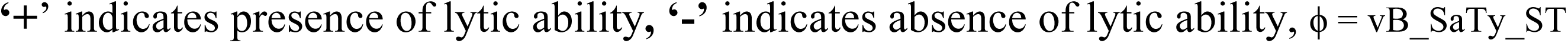
Host range determination of isolated phages.

| Bacterial<br>Strains | Bacteriophages |  |  |  |  |  |  |  |  |  |  |  |  |  |  |
| --- | --- | --- | --- | --- | --- | --- | --- | --- | --- | --- | --- | --- | --- | --- | --- |
| | ϕ1 | ϕ2 | ϕ3 | ϕ4 | ϕ5 | ϕ7 | ϕ8 | ϕ11 | ϕ12 | ϕ13 | ϕ15 | ϕ16 | ϕ17 | ϕ18 | ϕ19 |
| *S. Typhi* Ty2 | + | + | + | + | + | + | + | - | - | - | + | + | + | + | - |
| *S. Typhi* CT18 | - | - | - | - | - | - | - | + | + | + | - | - | - | - | + |
| *S. Typhimurium*<br>LT12 | - | - | - | - | - | - | - | - | - | - | - | - | - | - | - |
| *S. Typhi* ΔVi | - | - | - | - | - | - | - | - | - | - | - | - | - | - | - |
| *E. coli* 25922 | - | - | - | - | - | - | - | - | - | - | - | - | - | - | - |
| *S. aureus* 29213 | - | - | - | - | - | - | - | - | - | - | - | - | - | - | - |
| *E. faecalis* 51299 | - | - | - | - | - | - | - | - | - | - | - | - | - | - | - |
| *Klebsiella*<br>*quasipneumoniae*<br>700603 | - | - | - | - | - | - | - | - | - | - | - | - | - | - | - |
| *Pseudomonas*<br>*aeruginosa* 27853 | - | - | - | - | - | - | - | - | - | - | - | - | - | - | - |
| *E. faecalis* 29212 | - | - | - | - | - | - | - | - | - | - | - | - | - | - | - |
| *Klebsiella*<br>*pneumoniae* BAA-1705 | - | - | - | - | - | - | - | - | - | - | - | - | - | - | - |

‘+’ indicates presence of lytic ability, ‘-’ indicates absence of lytic ability, $\phi$ = vB\_SaTy\_ST

### Lytic ability against clinical isolates

Phages isolated against *S*. Typhi Ty2 were significantly more effective in lysing clinical *S*. Typhi isolates than those isolated against *S*. Typhi CT18 (odds ratio [OR] = 20.17, p <0.0001). Similarly, phage isolated against *S*. Typhi Ty2 were significantly more effective in lysing clinical *S*. Paratyphi isolates than those isolated against *S*. Typhi CT18 (OR = 1.68, p < 0.01) (Fig 5).

**Fig 5.**
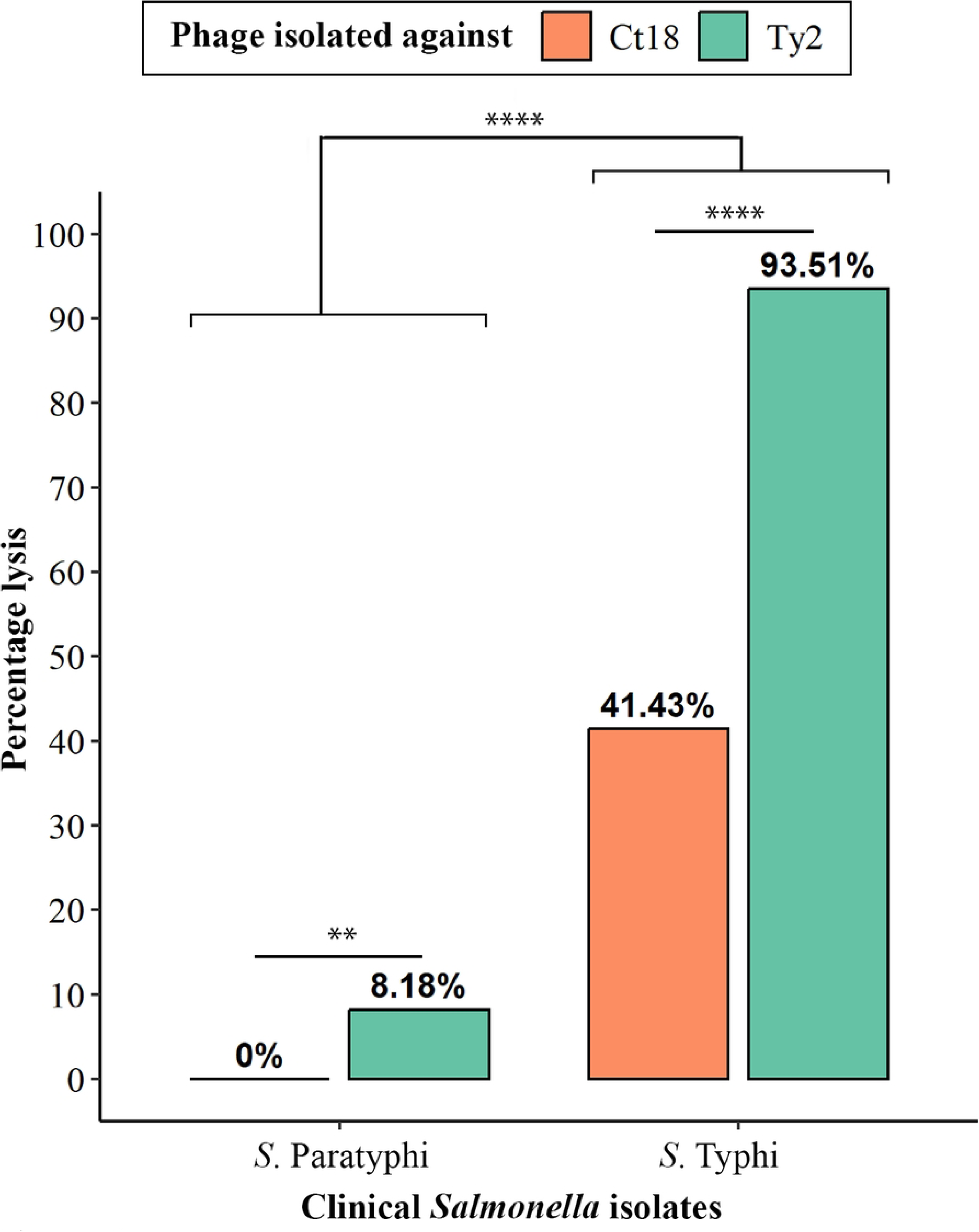
Lytic activity of bacteriophages against clinical *S*. Typhi and *S.* Paratyphi isolates. Bacteriophages isolated against S. Typhi Ct18 and S. Typhi Ty2 strains were tested for their ability to lyse clinical S. Paratyphi and S. Typhi isolates by Spot Assay. The statistical significance was tested using Fisher’s exact test; ** = p < 0.01, **** = p < 0.0001

Among the phages, phage vB_SaTy_ST2, vB_SaTy_ST7, vB_SaTy_ST17, and vB_SaTy_ST18 were the most effective to lyse 35 out of 35 (100%) *S*. Typhi isolates, while phage vB_SaTy_ST11 was the least effective, killing 7 out of 35 (20%) *S*. Typhi isolates (Fig 6). Out of all phages, only vB_SaTy_ST17 and vB_SaTy_ST18 could lyse 9 out of 20 (45%) *S*. Paratyphi isolates (S2 File).

**Fig 6.**
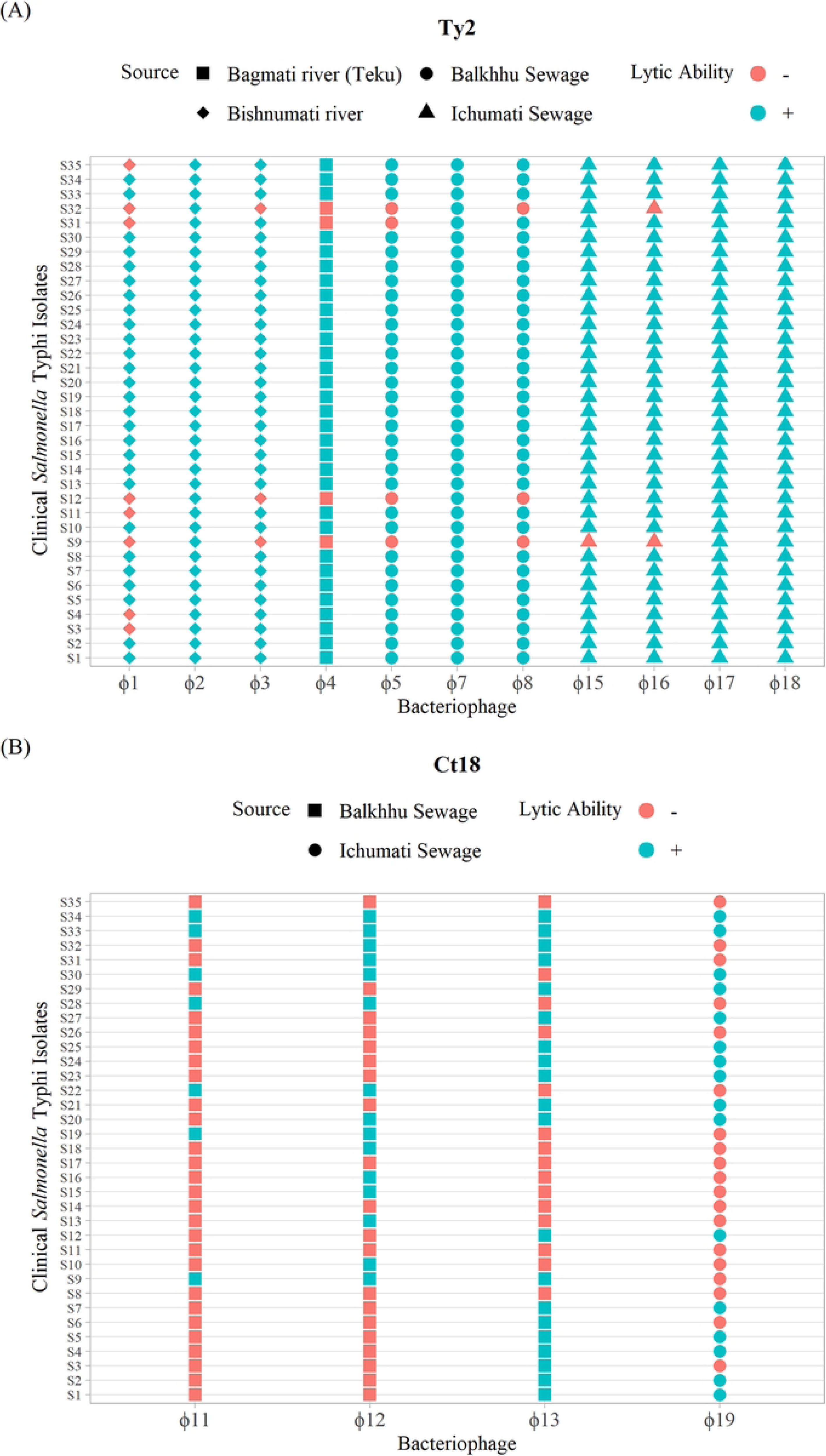
Lytic activity of individual bacteriophages against clinical *S*. Typhi isolates. The lytic ability of individual bacteriophages isolated against (A) S. Typhi Ty2 strain and (B) S. Typhi Ct18 strain was assessed on clinical isolates of S. Typhi using spot assay.; ϕ = vB_SaTy_ST

Overall, the isolated phages were significantly more effective at lysing clinical *S*. Typhi isolates than clinical *S*. Paratyphi isolates (OR = 60.69, p < 0.0001) (Fig 5).

### Effect of phage and phage cocktail on the growth kinetics of clinical *S.* Typhi isolate

The growth curves of *S.* Typhi isolate S30 infected by phage vB_SaTy_ST2, vB_SaTy_ST7, and vB_SaTy_ST17 at MOI adjusted to 0.1 showed individual phages were highly effective against *S*. Typhi S30. After 6 hours of incubation, *S.* Typhi S30 without phage (control) had reached the stationary phase with OD_600_ = 0.68, while no growth was detected when infected with individual phage. Phage vB_SaTy_ST17 demonstrated maximum efficacy in suppressing the growth of *S.* Typhi S30 up to 13 hours, while vB_SaTy_ST2 and vB_SaTy_ST7 independently could suppress the growth for up to 10 hours (Fig 7).

**Fig 7.**
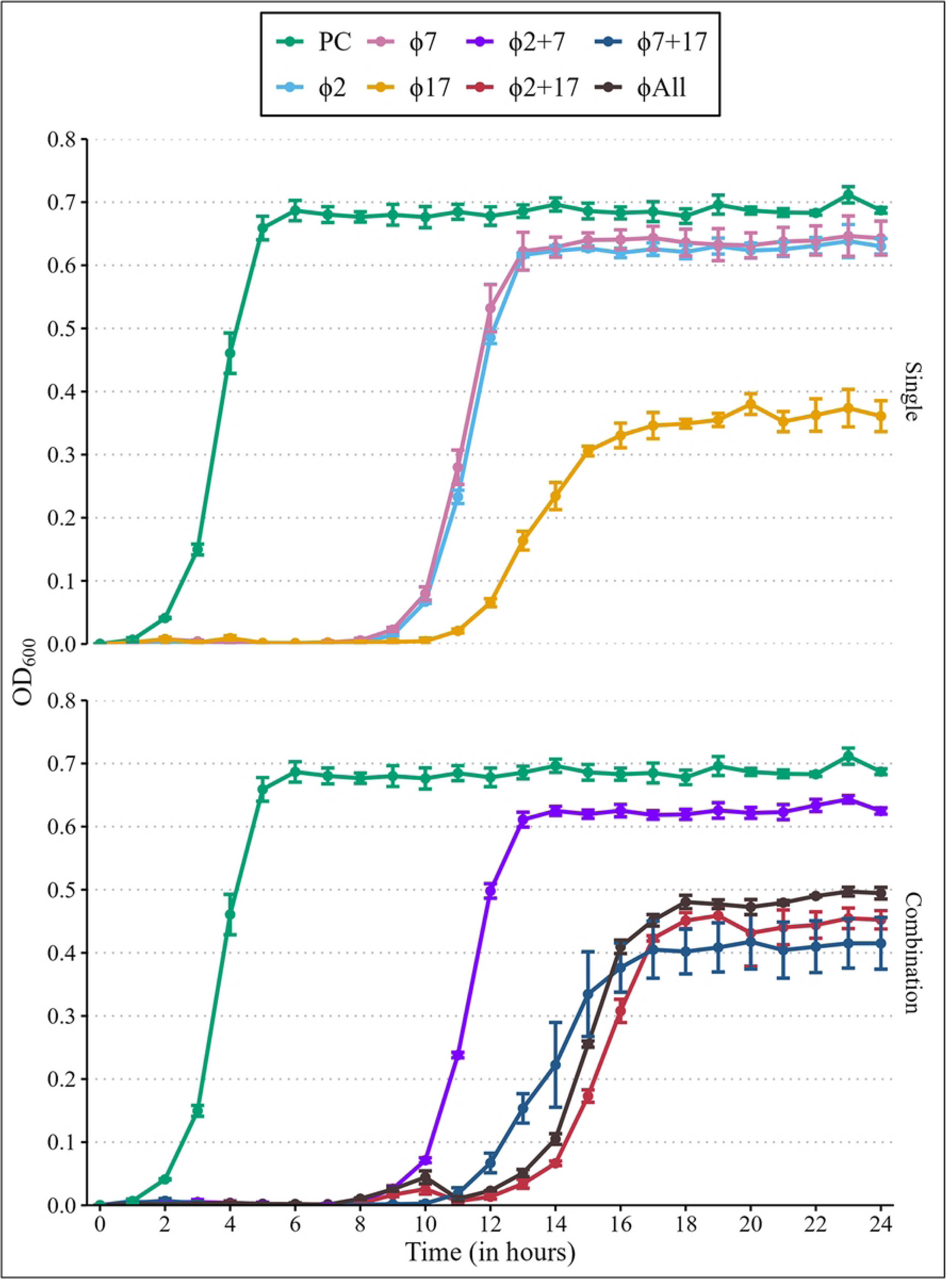
Effect of phage and phage cocktail on the growth of *S.* Typhi isolate S30. The in vitro growth kinetics were determined by the turbidimetric method. Sterile TSB was used as blank to measure the OD_600_ value, while *S*. Typhi S30 was used as a positive control (PC). OD_600_ value of 0.1 was considered an indicator of the early log phase.

The phage combination vB_SaTy_ST2 + vB_SaTy_ST17 inhibited the bacterial growth for up to 15 hours, while phage combination vB_SaTy_ST2 + vB_SaTy_ST7 and vB_SaTy_ST7 + vB_SaTy_ST17 inhibited bacterial growth up to 10 hours and 13 hours, respectively. The phage cocktail combining all three phages inhibited bacterial growth for up to 14 hours (Fig 7).

At the end of the experiment, the bacteria reached the stationary phase for all infections with individual phages and for all combinations of phage cocktails. However, all individual phages as well as all combinations of phages lowered the OD_600_ value of bacteria to reach the stationary phase as compared to bacteria grown alone (OD_600_ = 0.68). The OD_600_ reading was the lowest for bacteria infected with vB_SaTy_ST17 alone (OD600 = 0.36) (Fig 7).

## Discussion

Typhoid fever is endemic in Kathmandu Valley with seasonal incidence, which typically peaks during June to September [21,22]. In this study, the prevalence of enteric fever was found to be 3.32%. The rate of enteric fever was reported to be higher in 2012 (13.38%) at Kathmandu Model Hospital [23], which declined to 6.94% in 2018 from the same hospital [24] and shows a rather decreasing trend. A 23-year retrospective study conducted from 1992 to 2014 in Kathmandu Valley also highlighted the declining trend in overall enteric fever cases [22]. This decrease may be attributed to the increased access to safe drinking water and health facilities. Nonetheless, a study in 2021 reported a similar burden (3.1%) for enteric fever from a different hospital in Kathmandu [25]. In our study, *S*. Typhi was responsible for all enteric fever cases, which followed previous studies that reported the predominance of *S*. Typhi over *S*. Paratyphi [21,25], but it is in contrast to the finding that reported an increasing trend of *S*. Paratyphi infections [22].

In our study, enteric fever was more prevalent in young adults aged 16-25 years (48.57%), and the number of cases drastically decreased in adults and elderly people, which was similar to the previous finding, which reported 51% of enteric fever cases in the age group 16-25 years from Nepal [10]. Interestingly, enteric fever was not observed in the age group <5 years, which is unprecedented in previous studies [10,24,25]. This shift may be attributed to the introduction of typhoid conjugate vaccines (TCV) into Nepal’s routine immunization program in 2021 [26].

*S*. Typhi isolates in this study were sensitive to 1^st^ line drugs- chloramphenicol, ampicillin, and cotrimoxazole, i.e., none of the isolates were multidrug resistant. This finding is similar to previous studies that showed the re-emergence of conventional first-line antibiotics [23]. Interestingly, our study reveals a marked increase in the susceptibility of *S.* Typhi isolates to nalidixic acid and fluoroquinolones. While previous studies reported high levels of resistance to nalidixic acid (up to 96%) [27,28], our findings indicate an 80% susceptibility rate. This notable shift in antimicrobial susceptibility may be attributed to the introduction of the TCV. Earlier reports documenting high resistance were conducted during the pre-vaccination period, whereas our study is among the first to assess susceptibility patterns in the post-vaccination era.

In the current study, *S*. Typhi-specific bacteriophages were isolated from river water as well as sewage water. This shows that river and sewage water are excellent sources for phage isolation. Furthermore, phages were isolated more frequently from sewage water than from river water, suggesting sewage is a better source for phage isolation. Previous studies have also reported *S*. Typhi-specific phages to be more abundant in sewage-contaminated water in typhoid-endemic regions [29]. Sewage and sewage-contaminated water have a significant increase in microbial diversity due to mixing with human waste, which could explain the high number of phages in sewage [30].

The host range of a phage determines its utility in phage therapy. Ideally, phages should exhibit a narrow host range limited to the target pathogen, which ensures that normal flora is not affected by the therapy [31]. The isolated phages in this study were highly specific, with the ability to infect only *S.* Typhi. Most studies have reported *Salmonella*-specific phage to have a narrow host range [32,33] but there have been reports of polyvalent phage capable of infecting typhoidal as well as nontyphoidal strains [34]. The isolated phages could not infect the Vi polysaccharide knock-out *S*. Typhi ΔVi strain, indicating the need for Vi polysaccharide for infection.

Phage exhibiting broad lytic activity against multiple strains within the targeted bacterial species is highly sought after to be used in infections caused by that bacterium [31]. In our study, the isolated bacteriophages were highly effective in lysing clinical *S*. Typhi isolates with broad lytic activity; some were even capable of lysing all of the tested clinical *S*. Typhi isolates. The phages were highly specific and could discriminate between the typhoidal strains- *S*.

Paratyphi and *S*. Typhi, with the majority of phages capable of lysing only *S*. Typhi. This further enhances the potential of these phages as a therapeutic agents specifically targeting *S*. Typhi, the predominant typhoidal strain.

In our study, the isolated phages exhibited excellent bacteriolytic activity against the clinical isolate *S*. Typhi S30. The bacteria treated with phage vB_SaTy_ST2 and vB_SaTy_ST7 showed similar shifts in the growth curve, indicating these two phages might have similar replication strategies. The bacteria treated with phage vB_SaTy_ST17, however, had the most prominent inhibition in the growth rate. A similar study reported that phage vB_SalP_TR2 could inhibit the dynamic growth of *Salmonella* Albany at varying MOI [20]. Similarly, phage vB_StyS-LmqsSP1 was able to inhibit bacterial growth of *Salmonella* LT2 as well as two field isolates at MOIs as low as 0.0001 [35]. The bacterial growth dynamic is affected by multiple factors, including phage species, bacterial strain, and MOI, with infecting phage being the most important [36], which explains the different growth curve patterns obtained when treated with different phages.

Phage cocktails are developed mainly for two purposes: to increase the spectrum of targeted bacteria and to combat the potential for the evolution of phage-resistant bacteria [37]. In our study, the bacteriolytic effect of the phage cocktail consisting of phage vB_SaTy_ST2 + vB_SaTy_ST7 was not different from individual phages, which could suggest they have similar infection strategies. A synergistic effect was observed for phage cocktails vB_SaTy_ST2 + vB_SaTy_ST17 and vB_SaTy_ST2 + vB_SaTy_ST7 + vB_SaTy_ST17, resulting in greater inhibition of bacterial growth than treatment with individual phage. Our study showed phage cocktails can have a synergistic or no change in bacteriolytic effect, suggesting much consideration is required for selecting phages for phage cocktails. Phages with different host receptors are more suitable for phage cocktail preparation to combat phage-resistant bacteria when bacteria gain resistance by an adsorption blocking mechanism [38]. A systematically designed phage cocktail containing 5 phages targeting four receptors: O-antigen, BtuB, OmpC, and rough *Salmonella* strains was successful in achieving low emergence of resistance in *Salmonella enteritidis* [39]. The results from phage cocktail efficacy suggest phage vB_SaTy_ST2 and vB_SaTy_ST17 could have different host receptors, which is also supported by the observation that vB_SaTy_ST17 could infect *S*. Typhi as well as *S*. Paratyphi, while the lytic activity of vB_SaTy_ST2 is limited to only *S*. Typhi. Thus, vB_SaTy_ST2 + vB_SaTy_ST17 could be considered an effective cocktail to combat *S*. Typhi infections.

## Conclusions

This study revealed a considerable burden of enteric fever, with *S*. Typhi being the etiological agent for all infections. All *S.* Typhi isolates remained sensitive to conventional first-line drugs. Interestingly, this study reports the increase in susceptibility of *S*. Typhi to nalidixic acid and fluoroquinolones. This study also elucidated the isolation of *S*. Typhi-specific bacteriophages with remarkable lytic activity against clinical *S.* Typhi isolates. Three phages (vB_SaTy_ST2, vB_SaTy_ST7, vB_SaTy_ST17) with broad lytic activity were identified as potential therapeutic agents.

## Acknowledgments

This work was carried out with the aid of a grant from UENSCO-TWAS and Swedish International Development Cooperation Agency (Sida) grant no. 22-026 RG/BIO-CHE/AS_CG-FR 3240324952 to JS. The views expressed herein do not necessarily represent those of UNESCO-TWAS, Sida or its Board of Governors. We are grateful to the staff and faculty members of Kathmandu Model Hospital and the Institute for Research in Science and Technology for their support and coordination in accomplishing the study. We also thank all the patients for their involvement in the study.

## Supporting Information

**S1 Table. Plaque morphology of isolated phages**

**S2 Table. Lytic activity of individual bacteriophages against clinical S. Paratyphi isolates**

